# Population dynamics of *Cymodocea nodosa* under future ocean scenarios

**DOI:** 10.1101/484634

**Authors:** A.K. Mishra

## Abstract

Rising carbon dioxide (CO_2_) concentrations in the atmosphere will increase the average *p*CO_2_ level in the world oceans, which will have a knock-on effect on the marine ecosystem. Coastal seagrass communities are predicted to benefit from the increase in CO_2_ levels, but long-term effects of elevated CO_2_ on seagrass communities are less understood. Population reconstruction techniques were used to investigate the population dynamics of *Cymodocea nodosa* meadows, exposed to long term elevated CO_2_ at volcanic seeps off Greece and Italy. Effect of elevated CO_2_ was noticed on the growth, morphometry, density, biomass and age structure at CO_2_ seeps than reference sites. Above to below ground biomass ratio of *C. nodosa* were higher at CO_2_ seeps. The shoot age and shoot longevity of plants were lower at seeps. The present recruitment (sampled year) of the seagrass were higher than long-term average recruitment of the communities near the seeps. Carbon to nitrogen ratios (%DW) and annual leaf production of *C. nodosa* were higher in leaves at seeps. This study suggests under long-term CO_2_ enrichment C. *nodosa* production increases, but the plant survival rate decreases because of other co-factors such as nutrient availability and trace metal toxicity. Therefore, along with high CO_2_ other factors must be taken into consideration while predicting effects of future CO_2_ concentrations.

## Introduction

The ocean’s absorption of anthropogenic CO_2_ emissions (over 35 Giga tonnes of CO_2_ per year, (IPCC, 2014) has already lowered the mean ocean surface pH by 0.1 units since preindustrial values, with a predicted further decrease of 0.3-0.4 units by the end of this century (IPCC, 2014). These on-going changes are expected to intensify in the future with potentially significant, but variable effects on marine organisms depending on their sensitivity (Hendrick et al., 2010; Kroeker et al., 2010). Calcifying organisms are more susceptible to ocean acidification than non-calcifying organisms (Suggett et al., 2012), even though their responses to ocean acidification are dependent on the taxonomic group and their developmental stages (Hendrick et al., 2010; Kroeker et al., 2010). Realisation of the key role of seagrass in coastal ecosystems has fostered ever growing efforts to quantify their annual productivity and growth dynamics. (Duarte et al., 1994). Seagrasses, as many macro algal species, are notably tolerant to CO_2_ increases and may even benefit from it (Koch et al., 2013). In the current CO_2_ levels, seagrasses are dissolved inorganic carbon (DIC) limited as they are inefficient in utilising bicarbonate (Invers et al., 2001).

Hence, under increased CO_2_ conditions, it is expected seagrasses will increase utilization of CO_2_ (Beer et al. 1996) resulting in increased photosynthesis and consequently growth and productivity (Koch et al., 2013, Russel et al., 2013). Despite these predictions, the real-time studies conducted overtime both ex-situ with high CO_2_ and in-situ at natural CO_2_ seeps on seagrasses not completely support this expectation. For example, the photosynthetic activity of *Cymodocea nodosa* was stimulated by the low pH at natural CO_2_ seeps of Vulcano, with significant increase in chlorophyll-a content of leaves, maximum electron transport rate and compensation irradiance (Apostolaki et al., 2014). Low pH promoted productivity, but was not translated into biomass production, probably due to nutrient limitation, grazing or poor environmental conditions (Apostolaki et al., 2014). Similar results were obtained by Alexandra et al. (2012) for *C. nodosa* grown for five months under high CO_2_ conditions. On the other hand, the biomass and net primary production of *Halophila ovalis* and *Cymodocea serrulata* increased near the CO_2_ seeps, whereas the abundance of species increased only for *C. serrulata*, suggesting species specific response to elevated CO_2_ (Russell et al., 2013). Long term elevated CO_2_ experiments on *Zostera marina* for over one year showed greater reproductive outputs, increased below ground biomass and shoot density (Palacios and Zimmerman, 2007), whereas short term experiments resulted in increased photosynthetic rate and shoot productivity (Zimmerman et al., 1997). In contrast, experiments on *Cymodocea serrulata* have shown no enhancement in productivity at higher CO_2_ as they are carbon saturated in current CO_2_ concentrations (Schwarz et al., 2000). Recent CO_2_ enrichment studies on three tropical seagrass species showed a significant increase in net productivity with increase in CO_2_ levels, but a different growth rate between species was noticed due to varying strategies of carbon allocation among species (Ow et al., 2015).

Studies on natural CO_2_ seeps suggest that seagrass species can be adapted to survive and live under elevated CO_2_ conditions (Hall Spencer et al., 2008, Fabricius et al., 2011 and Takahashi et al., 2016). Much of the natural CO_2_ seeps of Europe are concentrated in the Mediterranean Sea (Dando et al.,1999). In the Mediterranean Sea, most known seeps are concentrated in the shallow waters of the active volcanic arcs in Aegean Seas and are usually of the gas hydrothermal type due to the large volume of gas released (Dando et al., 1999). These natural CO_2_ seeps provide future oceanic conditions (Hall Spencer et al.,2008; Hall-Spencer and Rodolfo-Metalpa, 2009) and are expected to affect seagrass communities due to changes in the physical and chemical features of seawater and sediments with possibly large effects on functioning features (Vizzini et al., 2010). In these conditions the growth and age structure of the seagrass *Cymodocea nodosa* in these seeps have not been investigated. The seagrass *Cymodocea nodosa* is an endemic species that supports highly complex and biodiverse climax communities in the Mediterranean Sea (Mazzella et al., 1986). Nevertheless, the effects of hydrothermal CO_2_ gas release associated to explosive volcanism activity on seagrass productivity has been studied (Vizzini et al., 2010). How these changes will affect higher levels of biological organization, such as seagrass population dynamics (e.g. shoot recruitment rate), is less studied.

The objective of this work is to assess for the first time the long-term responses of the population dynamics and production of seagrass, *C. nodosa* exposed to elevated CO_2_ levels. Populations in the vicinity and away from the influence of volcanic seeps were compared. CO_2_ seep sites have been used to assess the long-term effects of elevated CO_2_ on benthic marine ecosystems and respective underlying mechanisms (Hall-Spencer et al., 2008, Fabricius et al., 2011, Vizzini et al., 2013, Enochs et al., 2015). However, other cofounding factors related to the volcanic seeps, such as the emissions of heavy metals (Dando et al., 1999, Vizzini et al., 2013, Kadar et al., 2013) and sulphide (Dando et al., 1999, Boatta et al.,2013) may influence the plants and population responses to elevated CO_2_. These are very variable among seeps (Dekov et al., 2004, Varnavas et al., 2005) whereas the major CO_2_ composition of emissions is constant. To cope with possible confounding factors of the effects of CO_2_ on the population dynamics of *C. nodosa*, we replicated the sampling effort in three seeps, two at the island of Milos in Greece and one at Vulcano island in Italy, to consider only the common responses as effects of elevated CO_2_.

## Methods

### Study sites

#### Milos Islands, Greece

Paleochori Bay (36.67 N, 24.51 E) and Adamas thermal stations (36.70 N, 24.46 E) are part of Milos island (Fig.1A). Extensive submarine venting occurs offshore, from the intertidal to depths of more than 100 m over a 34 km^2^ area of seabed (Dando et al.,1999). The released gases are 95% CO_2_ with some H_2_S, CH_4_ and H_2_ (Dando et al., 1999).

#### Vulcano, Italy

We sampled Levante Bay (38.4 N, 15.0 E) off Vulcano island (Fig. 1B). Some parts of this bay are well-suited for studies of the effects of increased CO_2_ levels (Boatta et al., 2013) despite areas with elevated H_2_S and metals (Vizzini et al., 2013). The main underwater gas seeps are located along southern and western shores of the bay at <1 m depth (Boatta et al., 2013). Total CO_2_ output is about 3.6 tonne d^−1^ (Inguaggiato et al., 2012), and the underwater gas emissions are 97-98% CO_2_ with 2.2% H_2_S close to the seeps, decreasing to less than 0.005% H_2_S towards the north-eastern part of the bay, where most ocean acidification research has been located (Capaccioni et al., 2001; Milazzo et al., 2014). There is a step gradient in carbonate chemistry with pH 5.65 at the main gas seeps increasing to pH 8.1, which is typical for present day Mediterranean surface seawater, at >350 m from the seeps (Boatta et al., 2013). *Cymodocea nodosa* were absent at the main vents.

**Fig. 1.**
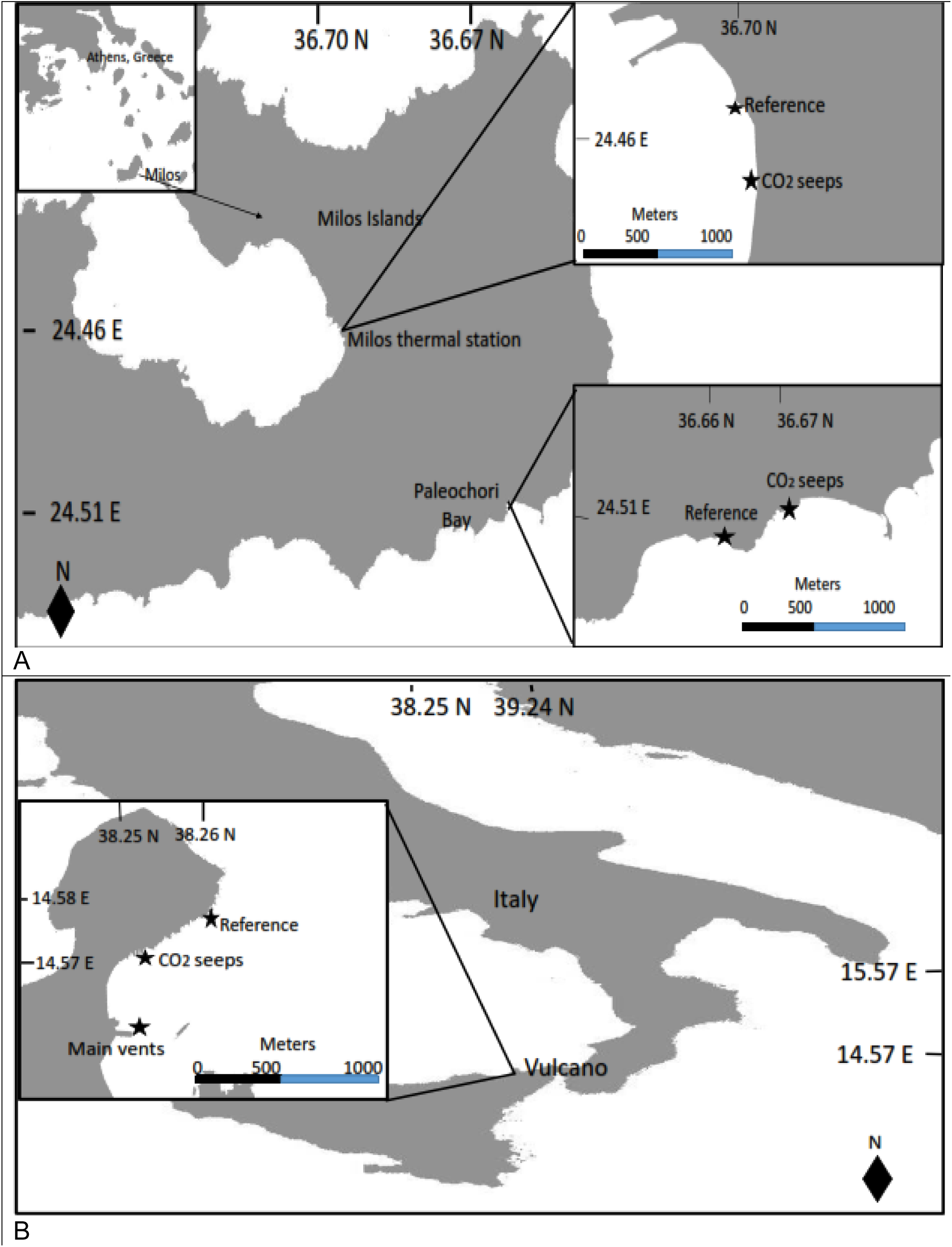
Study sites with location of CO_2_ seeps and Reference sites, A) Adamas and Paleochori off Greece and B) Vulcano island off Italy

In each location, two sites were established where well developed populations of *C. nodosa* were present, a high CO_2_ site near the seeps and a reference site away from the influence of the vent. Reference and CO_2_ seep sites were established at the same depths, under similar hydrodynamics conditions. Overall, *C. nodosa* meadows sampled were at shallow (<5m) depth.

#### Water sampling

Water samples (n=5) were collected at CO_2_ seeps and reference sites in 100 ml Winkler bottles fixed with 20 μl mercury chloride in the field, stored in the dark and transported to the laboratory for total alkalinity (TA) and dissolved inorganic carbon (DIC) analysis. The pH (using pH meter, Titrino Methron) and temperature of the water samples were measured in the field immediately after collection. In the laboratory pH and temperature were measured again and 80 ml of water sample was used in Lab Titrino analyser (Dickson et al. 2007) for the analysis of TA. Temperature, pH and TA data were used to calculate the other carbonate parameters (*p*CO_2_, HCO_3_^−^, aragonite saturation state, etc.) using CO_2_Sys program (Pierrot and Wallace.2006).

#### Seagrass sampling and analysis

The meadow density and biomass, the rhizome growth and production, the morphometric characteristics of the plants, the population age structure and derived population dynamics (long-term average recruitment, present recruitment and population growth rates) and the total C and N contents of plants were characterized in CO_2_ seeps and reference sites. Reconstruction techniques, an indirect measure of plant growth history and population dynamics changes (Duarte et al., 1994; Fourqurean et al. 2003), were used to evaluate the *C. nodosa* responses to the different environmental conditions caused by the seeps. It was hypothesized that increased CO_2_ promote higher plant growth, higher meadow production, and thus higher population dynamics.

Ten *C. nodosa* samples were collected from each site with a 20-cm diameter core (15 cm in Greece) into a depth of about 30 cm in May 2013 in Italy and May 2014 in Greece. The sediment was carefully rinsed off to prevent the modular sets disconnecting from each other and to keep the rhizome mat intact as required for the reconstruction of seagrass dynamics (Duarte et al., 1994). In each sample, the number of both shoots and apicals was counted to estimate the density of shoots and of physically independent individuals. The age of *C. nodosa* shoots was estimated by counting the number of leaf scars on the vertical rhizomes plus the number of leaves in each shoot multiplied by the leaf plastochrome interval (PI). To estimate the PI of each study site, i.e., the time needed to produce a new leaf, the sequence of average internodal length of *C. nodosa* shoots collected with the cores plus additional plants collected by hand was plotted. Then a 30% running average was applied to filter short-term seasonal variability and the difference in the number of vertical leaf scars between two consecutive length modes was counted. The modes represent annual growth periods and thus the average number of leaf scars produced between modes was averaged to estimate the leaf PI of the population (Short et al. 2001). The PI estimates of *Cymodocea nodosa* were 29.3 for all three locations off Greek and Italy islands.

To estimate the vertical and horizontal rhizome elongation rates the length of both the vertical and horizontal rhizomes between consecutive shoots was measured and the number of both vertical and horizontal internodes between consecutive shoots was counted (see Duarte et al., 1994 for details of method). The number of leaves per shoot were measured from intact shoots in each sample (n = 10). The horizontal and vertical rhizome production rates were estimated by multiplying the elongation rates (vertical or horizontal) by density (shoots or apicals), by the specific dry weight of rhizomes (vertical or horizontal) and by the dry weight contents of C (vertical or horizontal). Annual leaf production of each population was calculated as the product of the number of leaves annually produced per shoot, the shoot density, the mean specific dry weight of fully developed leaves and their C content.

The leaves, vertical rhizomes, horizontal rhizomes and roots were separated and dried for 48 h at 60° C for biomass and production estimates. Dried plant material (leaves, vertical rhizomes, horizontal rhizomes and roots) was grounded and analysed for CHN contents in a CHN analyser (EA 1110 Model, Elemental Microanalysis Ltd, Oakhampton, Devon, UK).

The long-term average recruitment (R) was estimated from the shoot age structure using the general model: N_x_ = N_0_ *e*^−Rx^, where N_x_ is the number of shoots in age class x, N_0_ is the number of shoots recruited into the population; assuming that mortality and recruitment have had no trend over the lifespan of the oldest shoots in the population, i.e. have remained constant over the lifespan of the oldest shoots, with year to year random variation around some mean value of mortality and recruitment (Fourqurean et al. 2003, Duarte et al. 2005). The recruitment for the current year of sampling (R_0_) was estimated using the method described by Duarte et al., (1994). The population growth rate (*r*) was estimated as: *r* = R_0_ – M, where M is the long-term mortality rate, which equals the long-term recruitment rate (R) under the assumptions of near steady state (Fourqurean et al., 2003). Population was considered growing if *r* is positive (R_0_ > R), shrinking if *r* is negative (R_0_ < R), or with the same trajectory pattern if R_0_ is not significantly different from R (Fourqurean et al., 2003).

#### Statistical analysis

Significant differences in biomass, density, production and plant morphometry among sites (CO_2_ seeps and reference) and locations (Adamas, Paleochori and Vulcano) were investigated using two-way ANOVA, after testing for homogeneity of variances and normality of distribution. The Tukey’s multiple comparison test was applied to determine significant differences between factor levels. When ANOVA assumptions were not verified, comparison of data sets were performed using the non-parametric test of Kruskal-Wallis and the post hoc Dunn’s test. Log transformations of variables were performed where needed.

The species vertical and horizontal rhizome elongation, and the population recruitment rates were obtained considering all replicates in each site. The t-test for the difference between two regression lines was used to compare the vertical rhizome elongation rates as these are equal to the slopes of the linear regression between age and size of rhizomes. Statistical analyses were not performed for the horizontal rhizome elongation rate, because just one value was obtained for each site. The confidence limits of the exponential decay regression model used to estimate the long-term average recruitment rate (R) allowed its statistical comparison to the present recruitment rate (R_0_) as described in Fourqurean et al., (2003). Significant differences of the long-term recruitment rate among sites were tested using one-way ANOVA. Significance levels was considered at *p* < 0.05 (Sokal and Rohlf, 2012).

## Results

Seawater carbonate chemistry values (mean ± se) of the three sites Adamas, Paleochori and Vulcano are presented in Table 1. At all stations, the *p*CO_2_ concentration was higher (737± 158 - 2457.9± 1.87) and pH lower (7.98± 0.08 - 7.50± 0.04) near the CO_2_ seeps (Table 1). The reference stations (Reference) lying further away from the CO_2_ seeps showed almost two times lower concentrations of CO_2_ except for Vulcano. DIC (3474.03± 4.55) and CO_2_ (2377± 37) concentrations were higher at Adamas and lower at Vulcano.

**Table 1.** Seawater carbonate chemistry measurements at Adamas, Paleochori and Vulcano CO_2_ seeps calculated with CO_2_Sys programme, using constants from Dickson and Millero, 1987, and pH in NBS scale.

|  | **CO<sub>2</sub> seeps** | **Reference** |
| --- | --- | --- |
|  | **Adamas** |  |
| pH (NBS scale) | 7.5 ± 0.04 | 8.2 ± 0.03 |
| *p*CO<sub>2</sub> (μ atm) | 2457.9 ± 1.87 | 405.5 ± 1.65 |
| DIC (μ mol kg<sup>-1</sup>) | 3474.03 ± 4.55 | 1407.44 ± 51.25 |
|  | **Paleochori** |  |
| pH (NBS scale) | 7.9 ± 0.01 | 8.2 ± 0.01 |
| *p*CO<sub>2</sub> (μ atm) | 884.3 ± 3.07 | 402.9 ± 1.1 |
| DIC (μ mol kg<sup>-1</sup>) | 2738.39 ± 16.49 | 1396.14 ± 48.69 |
|  | **Vulcano** |  |
| pH (NBS scale) | 7.98 ± 0.08 | 8.17 ± 0.05 |
| *p*CO<sub>2</sub> (μ atm) | 737 ± 158 | 427 ± 68 |
| DIC (μ mol kg<sup>-1</sup>) | 2377 ± 37 | 2244 ± 40 |

Significant response of *C. nodosa* to elevated CO_2_ was noticed on density and biomass at all three CO_2_ seeps. Both shoot and apex density, i.e. the number of physically independent plants was 1.3fold (shoot density) and 2.3fold (apex density) higher at CO_2_ seeps than reference stations, exceptions were shoot density at Vulcano and apex density at Adamas seep stations (Fig. 2). Both shoot (1322.00± 166.20 no.m^−2^) and apex density (299.40±37.70 no.m^−2^) was higher at Adamas than Paleochori and Vulcano seep stations. Total biomass (456.20 ± 64.90 g DW m^−2^), leaf biomass (97.70± 9.10 g DW m^−2^) and horizontal biomass (155.10± 29.0 g DW m^−2^) were higher at seeps of Vulcano than Adamas and Paleochori stations, whereas above ground-below ground biomass ratio (0.60± 0.10 g DW m^−2^) was higher at seeps of Paleochori (Fig. 2). Significant increase (>1.2-fold) in total biomass and leaf biomass was observed at Paleochori and Adamas seeps than reference stations, whereas for horizontal biomass this increase (>1.2-fold) was found at seeps of Paleochori and Vulcano. However, the highest total biomass recorded for Vulcano was 0.9-fold lower than reference station. The vertical rhizome biomass (67.60± 10.0 g DW m^−2^) was significantly lower near the seeps than reference (124.40± 17.60 g DW m^−2^) stations of Vulcano but significant increase of biomass was observed for Paleochori (1.9-fold) and Adamas (1.2-fold). Significant effects of CO_2_ on plant morphology was observed on number of leaves and vertical rhizome. The number of leaves per shoot was higher at Adamas (4.50± 0.01shoot^−1^) and the vertical rhizome (0.35 ± 0.03-2.10± 0.10 cm) was shorter near all the seep stations (Fig.3).

**Fig. 2.**
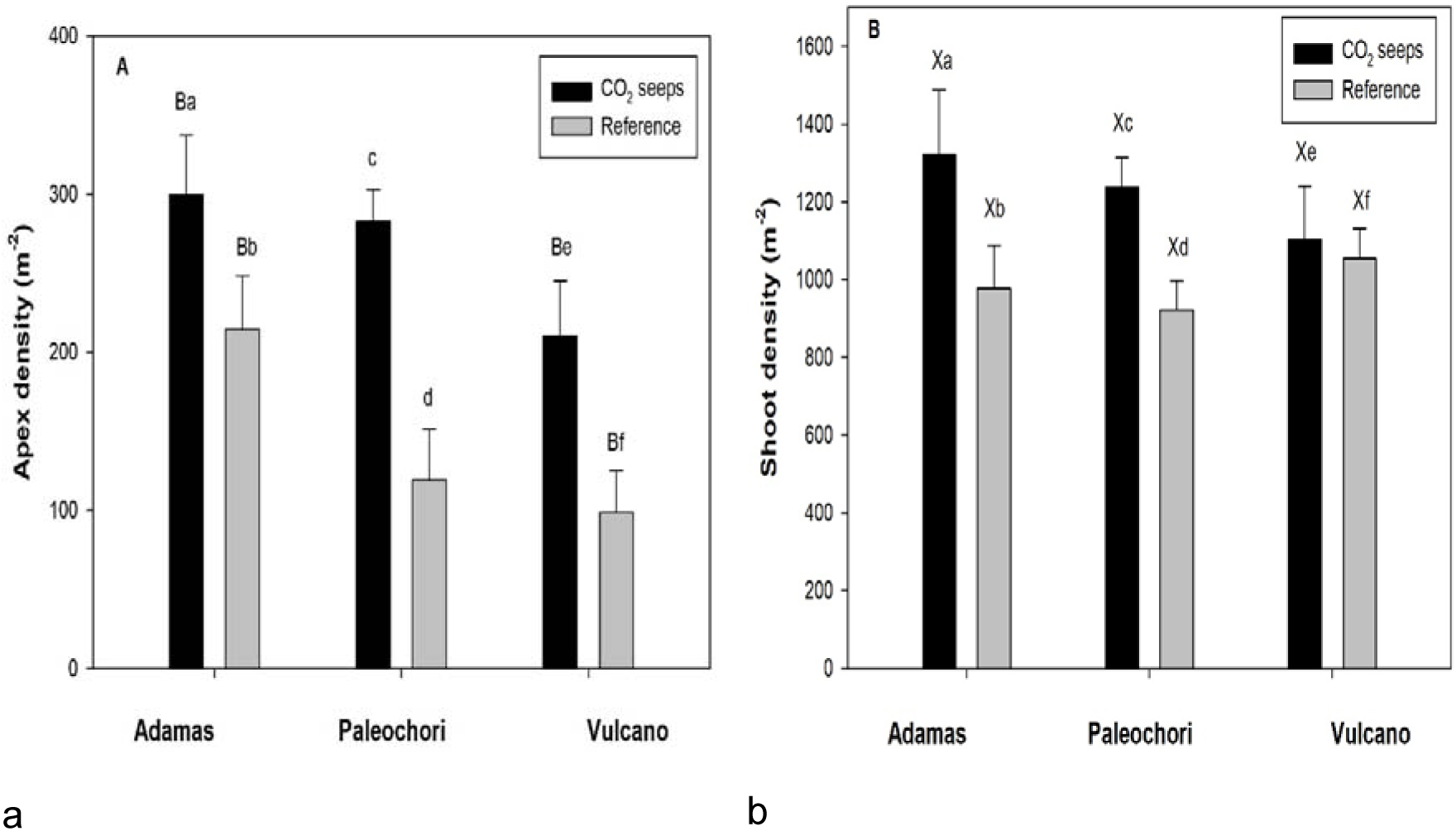
Density (Apex and Shoot) of *Cymodocea nodosa* at the CO_2_ seeps off the Greek (Adamas and Paleochori) and Italy (Vulcano) islands. Error bars represent standard errors. Differences between sites (Adamas vs Paleochori-A, Adamas vs Vulcano-B, Vulcano vs Paleochori-C) and stations (CO_2_ seeps and reference) are indicated by capital letters and small letters respectively.

**Fig. 3.**
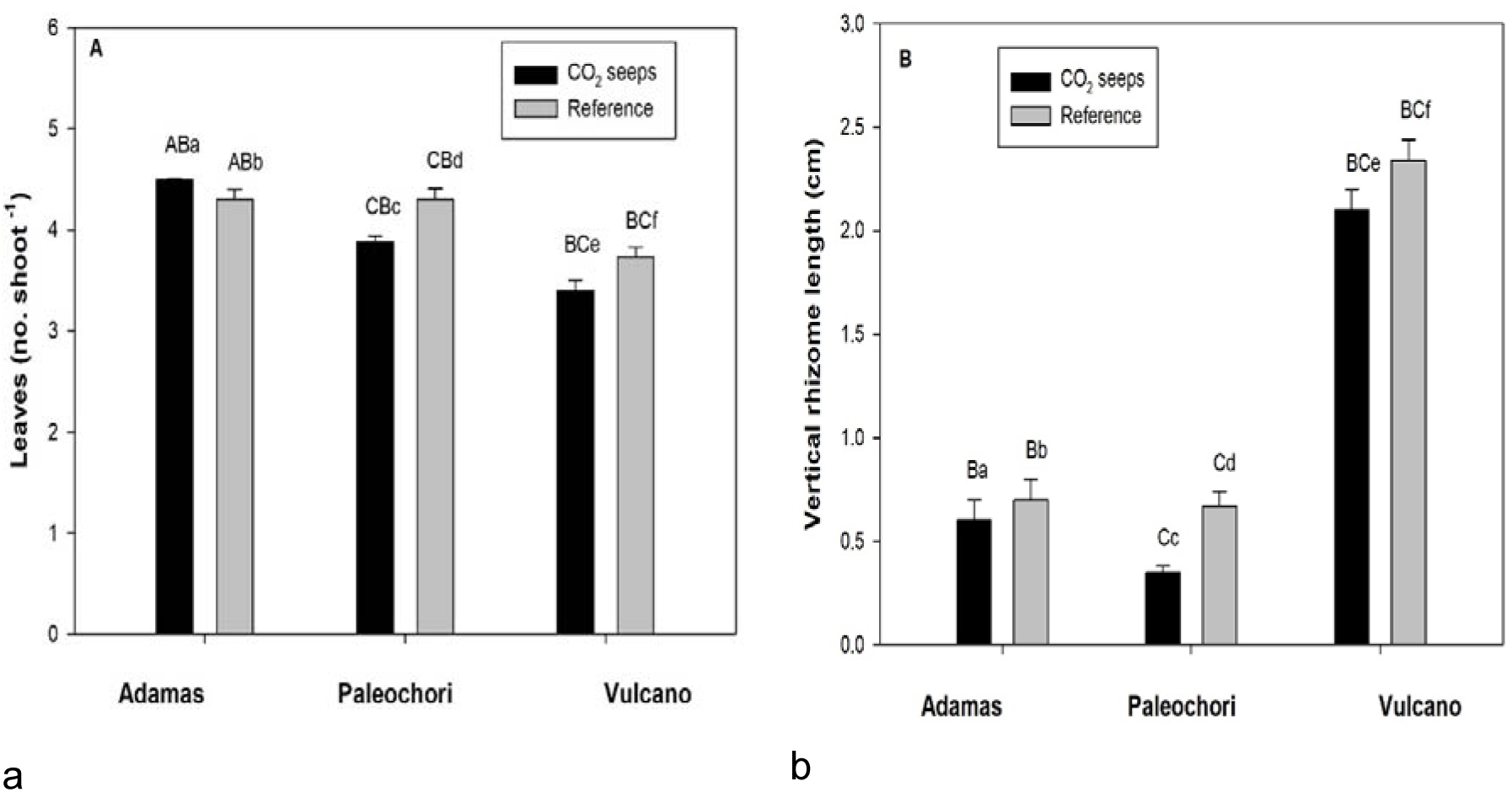
Morphometry (Number of Leaves and Vertical rhizome length) of *Cymodocea nodosa* at CO_2_ seeps and reference sites off Greek (Adamas and Paleochori) and Italy (Vulcano) islands. Error bars represent standard errors. Differences between sites (Adamas vs Paleochori-A, Adamas vs Vulcano-B, Vulcano vs Paleochori-C) and stations (CO_2_ seeps and reference) are indicated by capital letters and small letters respectively.

**Fig. 4.**
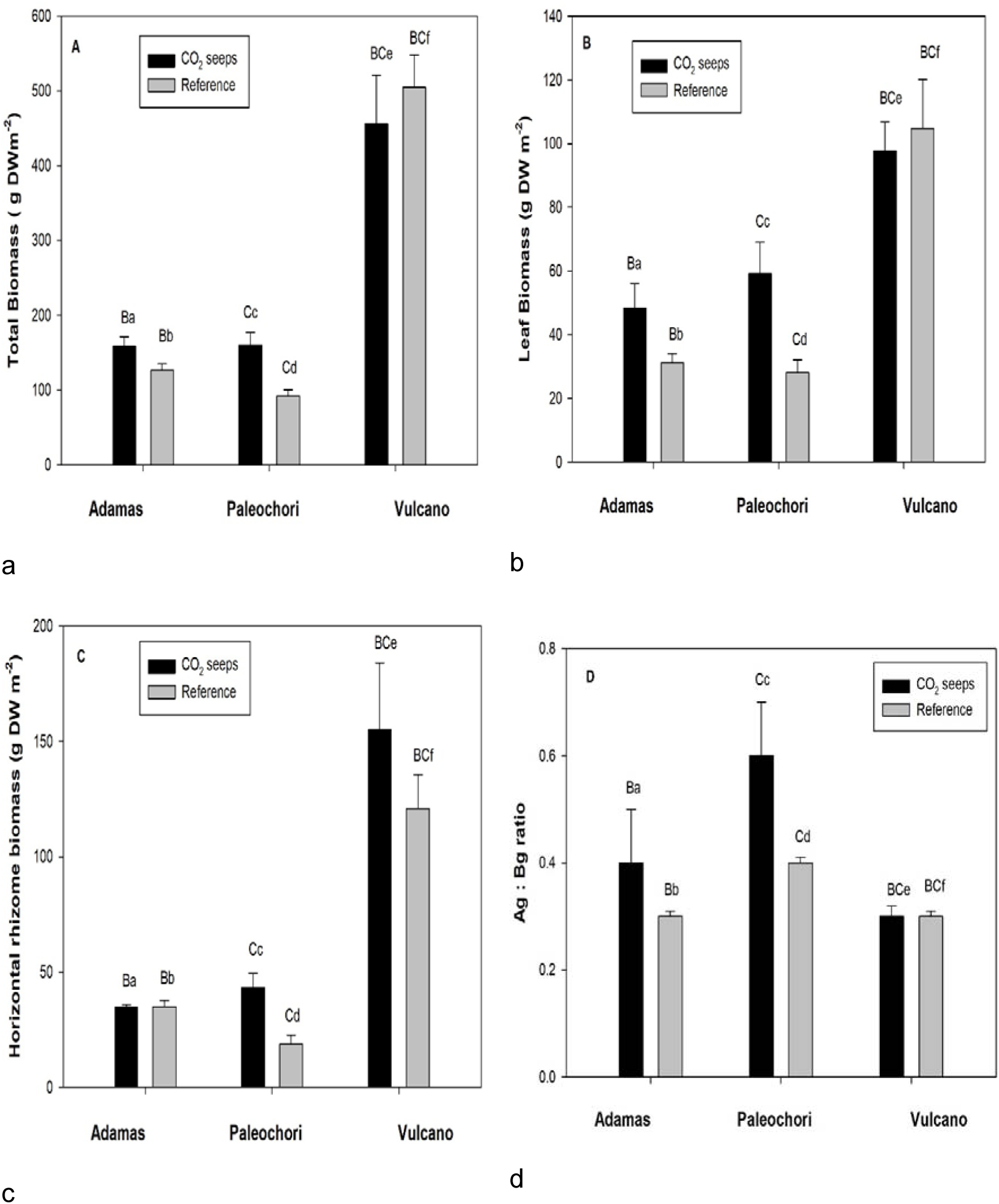
Biomass of *Cymodocea nodosa* at the CO_2_ seeps and reference sites off Greek (Adamas and Paleochori) and Italy (Vulcano) islands. Error bars represent standard errors. Differences between sites (Adamas vs Paleochori-A, Adamas vs Vulcano-B, Vulcano vs Paleochori-C) and stations (CO_2_ seeps and reference) are indicated by capital letters and small letters respectively (Ag: Bg; Above ground biomass: Below ground biomass)

Significant variation in density, biomass and morphometry of *C. nodosa* was observed between Greek and Italian sites (Adamas vs Vulcano, Paleochori vs Vulcano) but not between Greek sites (Adamas vs Paleochori), except for number of leaves which was significant between Greek sites and shoot density that did not vary among sites (Fig.3). Significant differences between apex density was found between Adamas vs Vulcano and for biomass between sites of Adamas vs Vulcano and Vulcano vs Paleochori. The vertical rhizome length and number of leaves varied significantly between Adamas vs Vulcano and Vulcano vs Paleochori (Fig.3).

Vertical elongation rate of *C. nodosa* was 1 to 1.2-fold higher at CO_2_ seeps than reference stations at all sites and significant (see Appendix). Horizontal elongation rate was1 to 1.3-fold higher at seep than reference stations at all sites except for Vulcano CO_2_ seep station, where it was 0.7-fold lower than reference station (See Appendix).

The elemental content of carbon and C: N ratios of *C. nodosa* leaves, rhizomes and roots showed that whenever there are significant differences, these reflect a carbon enrichment of tissues near CO_2_ seeps (Table 3). Significantly higher carbon content was observed near the seep in the leaves and roots at Adamas, leaves at Paleochori and leaves and rhizomes at Vulcano than reference stations. The only significant difference found in the tissues N content was in the rhizomes of Paleochori, where it was 4.2-fold lower near the seeps than reference station (Table 3). Significant differences in C: N ratios were observed in roots of Adamas, rhizomes of Paleochori and roots and rhizomes of Vulcano. Significant differences of C and N content in leaves among sites was found at Adamas vs Vulcano and Vulcano vs Paleochori. In the roots the C and N content was significant al all three sites (Table 3). The C: N content significant differences was found in the leaves and rhizomes at Adamas vs Paleochori and Vulcano vs Paleochori.

**Table 2.** Age structure and population dynamics of *Cymodocea nodosa* shoots at CO_2_ seeps and reference sites of Greek (Adamas and Paleochori) and Italy (Vulcano) islands. Mean ± Standard errors are presented for the shoot age. The exponential coefficient ± standard errors of the exponential decay regression are presented for the long-term average recruitment rate (R). Different letters indicate significant difference between CO_2_ seeps and reference sites only for R, ns= not significant. P values obtained from one-way ANOVA for Adamas (p=0.116), Paleochori (p=0.013) and Vulcano (p= 0.105)

|  | **Adamas** |  |
| --- | --- | --- |
|  | **CO<sub>2</sub> seeps** | **Reference** |
| Shoot longevity (years) | 3.21 | 3.53 |
| Long term avg. recruitment rate (R, year<sup>-1</sup>) | 1.03 ± 0.05<sup>ns</sup> | 0.90 ± 0.05<sup>ns</sup> |
| Present recruitment rate (R<sub>0</sub>, year<sup>-1</sup>) | 1.08 | 0.94 |
| Population growth rate (r, year<sup>-1</sup>) | 0.05 | 0.04 |
|  | **Paleochori** |  |
|  | **CO<sub>2</sub> seeps** | **Reference** |
| Shoot longevity (years) | 2.41 | 2.81 |
| Long term avg. recruitment rate (R, year<sup>-1</sup>) | 1.41 ± 0.08<sup>a</sup> | 0.42 ± 0.23<sup>b</sup> |
| Present recruitment rate (R<sub>0</sub>, year<sup>-1</sup>) | 1.54 | 0.52 |
| Population growth rate (r, year<sup>-1</sup>) | 0.07 | 0.09 |
|  | **Vulcano** |  |
|  | **CO<sub>2</sub> seeps** | **Reference** |
| Shoot longevity (years) | 7.14 | 8.27 |
| Long-term avg. recruitment rate (R, year<sup>-1</sup>) | 0.64 ± 0.11<sup>ns</sup> | 0.37 ± 0.05<sup>ns</sup> |
| Present recruitment rate (R<sub>0</sub>, year<sup>-1</sup>) | 0.65 | 0.36 |
| Population growth rate (r, year<sup>-1</sup>) | 0.01 | -0.01 |

The effects of elevated CO_2_ were also observed on *C. nodosa* rhizome and leaf production. The vertical rhizome production was 2.1 to 2.7-fold higher and horizontal rhizome production were 1.3 to 2.9-fold higher at CO_2_ seep than reference stations (Fig.5). Highest vertical (200.0± 60.20 gCm^−2^ y^−1^) and horizontal rhizome (299.50± 37.70 gCm^−2^ y^−1^) production was found at Adamas and lowest at Vulcano seep stations. Annual leaf production was 1.2-fold higher at seeps than reference stations with highest leaf production (245.04± 26.68 gCm^−2^ y^−1^) found at Vulcano seep station (Fig.6). Leaf production of Vulcano and Adamas was 2.3-fold and 1.1-fold higher than Paleochori seep station. The vertical rhizome production was significantly different at Vulcano vs Adamas and Adamas vs Paleochori sites and for horizontal rhizome at all three CO_2_ seep sites. Significant differences for leaf production were found for Adamas vs Vulcano and Vulcano vs Paleochori (Fig.6).

**Fig. 5.**
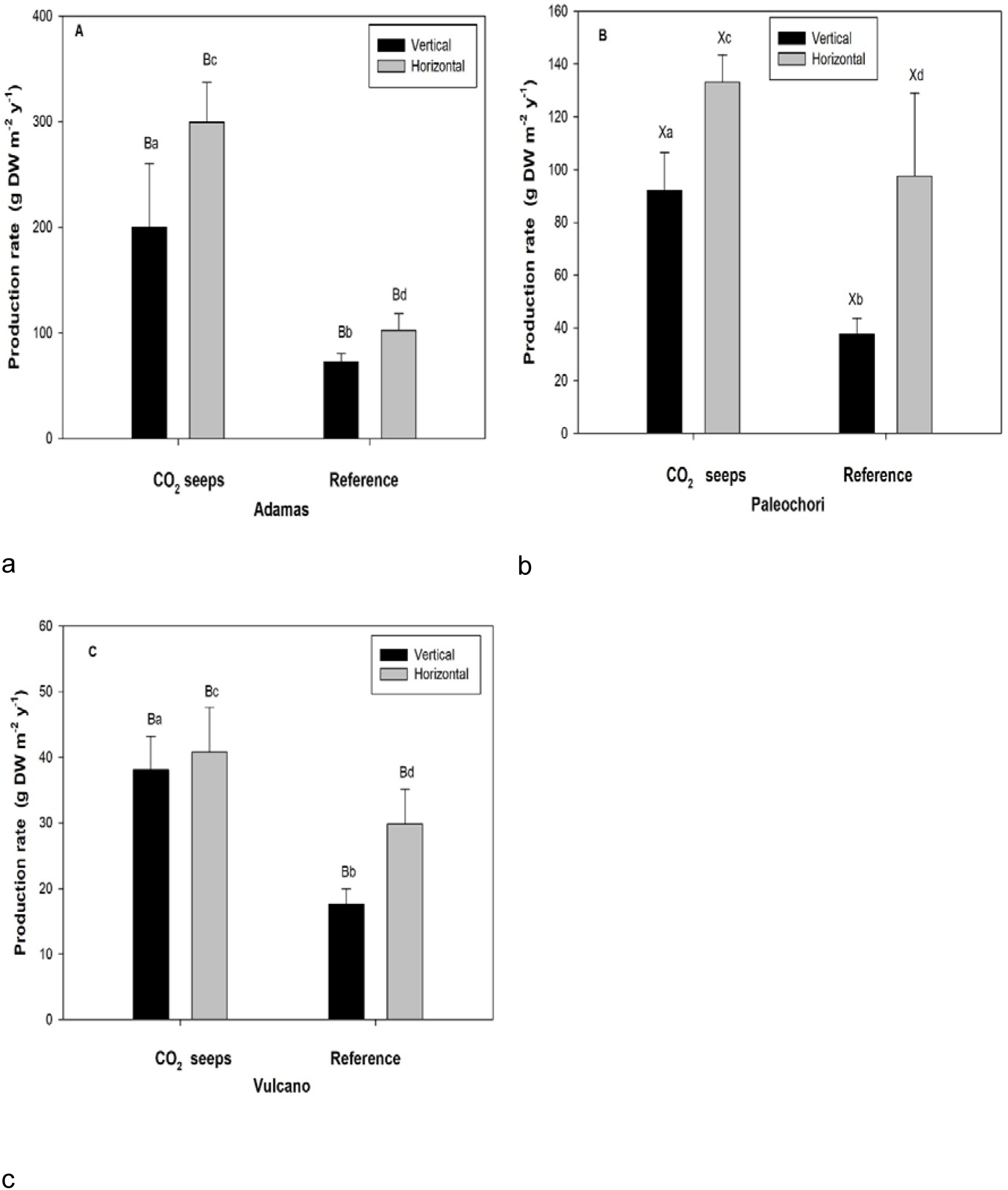
Vertical and horizontal rhizome productions of *Cymodocea nodosa* at the CO_2_ seeps and reference sites of Greek (Adamas and Paleochori) and Italy (Vulcano) islands. Error bars represent standard errors. Differences between sites (Adamas vs Paleochori-A, Adamas vs Vulcano-B, Vulcano vs Paleochori-C) and stations (CO_2_ seeps and reference) are indicated by capital letters and small letters respectively. No difference between sites are indicated by X.

**Fig. 6.**
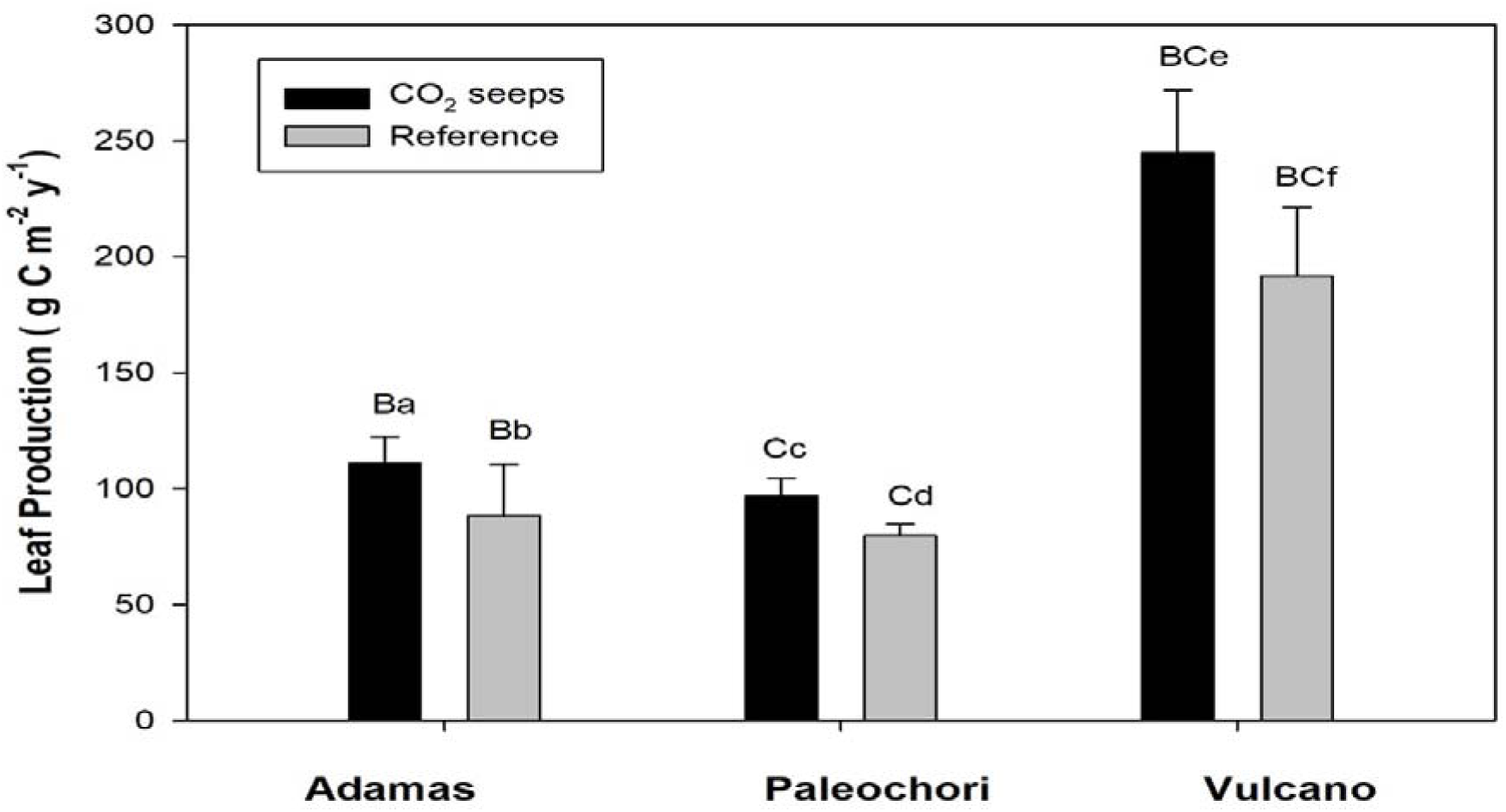
Annual leaf Production of *Cymodocea nodosa* at the CO_2_ seeps and reference sites off Greek (Adamas and Paleochori) and Italy (Vulcano) islands. Error bars represent standard errors. Differences between sites (Adamas vs Paleochori-A, Adamas vs Vulcano-B, Vulcano vs Paleochori-C) and stations (CO_2_ seeps and reference) are indicated by capital letters and small letters respectively.

The age structure of *C. nodosa* was significantly affected at CO_2_ seeps, as both the average shoot age (Fig.7) and shoot longevity, were lower near the seeps at all sites (Table 2). The lifespan of the oldest shoots was between seven to eight years at Vulcano, whereas at Paleochori and Adamas it varied between three to four years (Fig. 7). The present and long-term recruitments were always higher near the seeps, even though the differences were not statistically significant, except for Paleochori (Table 2). The age frequency distribution of *C. nodosa* showed a significant higher number of younger plants (1 and 2 years older) at the CO_2_ seeps, whereas the number of older plants (more than 3 years) were higher near the reference stations at all three sites (Fig.8).

**Fig. 7.**
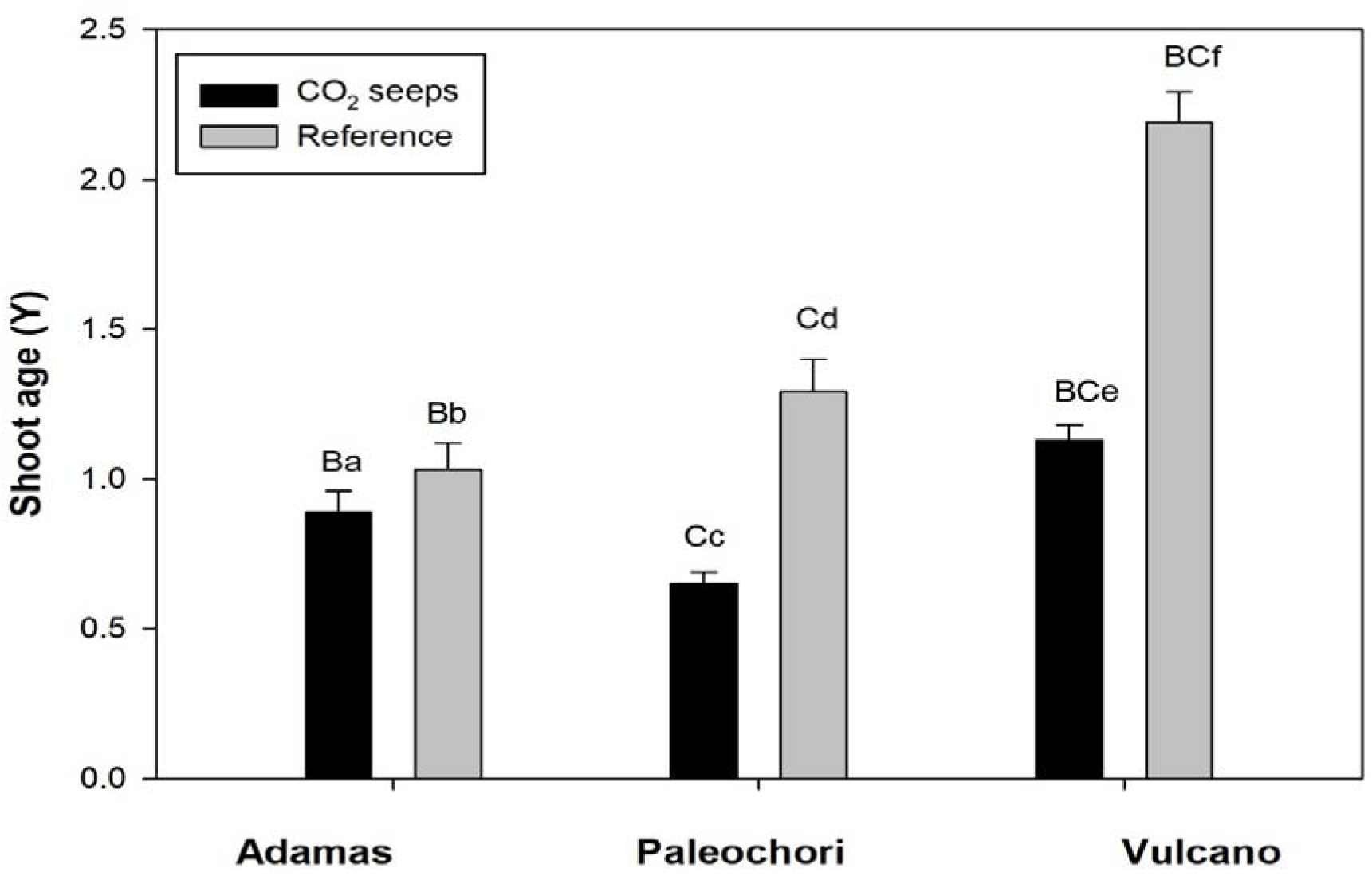
Shoot age of *Cymodocea nodosa* at the CO_2_ seeps and reference sites off Greek (Adamas and Paleochori) and Italy (Vulcano) islands. Error bars represent standard errors. Differences between sites (Adamas vs Paleochori-A, Adamas vs Vulcano-B, Vulcano vs Paleochori-C) and stations (CO_2_ seeps and reference) are indicated by capital letters and small letters respectively.

**Fig. 8.**
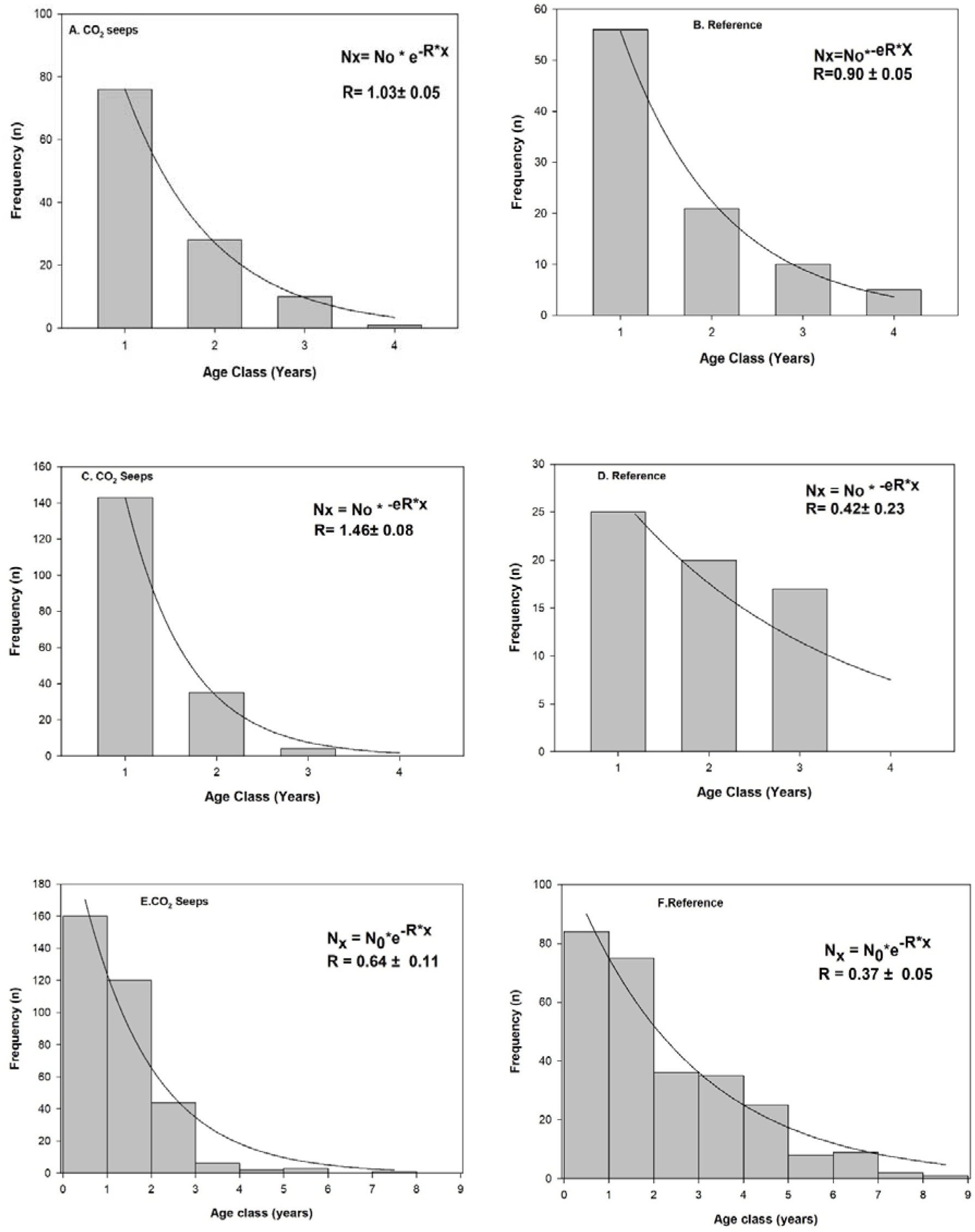
Age frequency distribution of *Cymodocea nodosa* population along the CO_2_ seeps and reference sites off Adamas (a, b), Paleochori (c, d) and Vulcano (e, f) respectively. The long-term average recruitment rate (R) was estimated from the exponential decay regression line fitted to age frequency distribution.

**Table 3.** *Cymodocea nodosa* carbon (C) and nitrogen (N) content and ratios (means ± SE) at CO_2_ seeps of Greek (Adamas and Paleochori) and Italy (Vulcano) islands. Tukey’s multiple comparison (p<0.05) test results are presented. Differences between sites (Adamas vs Paleochori-A, Adamas vs Vulcano-B, Vulcano vs Paleochori-C) and stations (CO_2_ seeps and reference) are indicated by capital letters and small letters respectively. No difference between sites are presented by X

| Variables |  | Adamas |  | Paleochori |  | Vulcano |  |
| --- | --- | --- | --- | --- | --- | --- | --- |
|  |  | CO<sub>2</sub> seeps | Reference | CO<sub>2</sub> seeps | Reference | CO<sub>2</sub> seeps | Reference |
| **C (%DW)** | Leaves | 35.0±0.19<sup>Ba</sup> | 31.9±0.70<sup>Bb</sup> | 35.0±0.80<sup>Cc</sup> | 35.2±0.51<sup>Cc</sup> | 41.9±0.62<sup>BCd</sup> | 41.3±0.03<sup>BCd</sup> |
| | Rhizomes | 34.6±0.23<sup>Xa</sup> | 32.9±0.70<sup>Xa</sup> | 31.7±0.63<sup>Xb</sup> | 33.1±0.38<sup>Xb</sup> | 33.4±0.37<sup>Xc</sup> | 33.0±0.4<sup>Xd</sup> |
| | Roots | 35.3±0.47<sup>ABa</sup> | 34.1±0.25<sup>ABa</sup> | 31.9±0.71<sup>Ab</sup> | 33.0±0.42<sup>BCb</sup> | 30.5±0.40<sup>BXc</sup> | 28.8±0.6<sup>CXc</sup> |
| **N (%DW)** | Leaves | 1.8±0.05<sup>Ba</sup> | 1.7±0.01<sup>Ba</sup> | 1.6±0.36<sup>Cb</sup> | 1.4±0.08<sup>Cb</sup> | 2.5±0.01<sup>BCc</sup> | 2.3±0.4<sup>BCc</sup> |
| | Roots | 0.9±0.17<sup>Xa</sup> | 0.9±0.11<sup>Xa</sup> | 0.8±0.11<sup>Xb</sup> | 1.0±0.07<sup>Xb</sup> | 0.8±0.02<sup>Xc</sup> | 1.0±0.1<sup>Xc</sup> |
| **C: N** | Leaves | 18.7±0.12<sup>Aa</sup> | 18.1±0.36<sup>Ab</sup> | 20.9±0.80<sup>ACc</sup> | 15.3±0.37<sup>Ad</sup> | 16.4± 0.28<sup>Ce</sup> | 17.5±0.6<sup>Ce</sup> |
| | Rhizomes | 20.5±0.24<sup>Aa</sup> | 23.5±0.85<sup>Aa</sup> | 34.2±0.63<sup>ACb</sup> | 33.0±0.42<sup>Xc</sup> | 23.5±1.68<sup>Cc</sup> | 26.3±1.0<sup>Xd</sup> |
| | Roots | 37.5±0.32<sup>Xa</sup> | 36.3±0.18<sup>Xb</sup> | 39.7±0.71<sup>Xc</sup> | 17.5±0.62<sup>Cd</sup> | 34.1± 1.60<sup>Xe</sup> | 28.9±3.37<sup>Xe</sup> |

The different variables analysed here for *C. nodosa* using reconstruction techniques responded differently between CO_2_ seep and reference stations among the three sites of Adamas, Paleochori and Vulcano of Greek and Italy islands. Significant interaction among stations and sites for variables (number of leaves, total biomass, horizontal rhizome biomass, above ground: below ground biomass, Leaves (%C), Roots (%C), Rhizomes (%N), Rhizomes (C: N)) was observed at all three sites of Adamas, Paleochori and Vulcano.

## Discussion

Population dynamics of *C. nodosa* near the shallow CO_2_ seeps of Greek (Adamas and Paleochori) and Italy(Vulcano) islands were positively affected under long term increased CO_2_ levels. The density, biomass, morphometry, age structure and production of *C. nodosa* showed positive response to elevated CO_2_ levels, although this response was variable among the three sites.

The high *p*CO_2_ level and low pH at the CO_2_ seeps of the Greek and Italy islands represented current (reference) and future 2100 (elevated CO_2_) scenarios, enabling us to assess the long-term effects of elevated CO_2_ on seagrass communities. Input of volcanic CO_2_ concentrations from the seeps, not only increased the dissolved CO_2_ concentration in the surrounding waters, but also increased the relative portion of dissolved CO_2_ to HCO_3_- (Short and Neckles. 1999) with a positive effect on seagrasses, as observed for the vertical and horizontal rhizome productivity, apex and shoot density and biomass of *C. nodosa* found at all seeps investigated. Plants developing near the CO_2_-seeps expressed an increase in fitness confirming that seagrasses can thrive in predicted scenarios of global CO_2_/OA changes (Hall-Spencer et al. 2008; Russell et al. 2013). Boosted growth and production with proximity to CO_2_-seeps suggest that the increase in CO_2_ concentrations will favour the utilization of inorganic carbon sources, seagrasses being C limited (Invers et al., 2001; Apostolaki et al., 2014)

The distribution of *C. nodosa* around the seeps may be related to high level of patchiness of the seagrass meadow, which was evident during sampling. Apex density of *C. nodosa* were higher at all three CO_2_ seeps compared to reference sites, but the higher density was not transferred to significant patch expansion of seagrass meadows close to the seeps, even though *C. nodosa* can translocate itself to make meadows (Kraemer and Mazzella, 1999). Lack of *C. nodosa* meadows near the CO_2_ seeps can also be related to the associated geochemical features (e.g. toxic levels of trace elements) of individual CO_2_ seeps affecting the migration of meadow spatially. Increase in density at all three CO_2_ seeps and biomass (only at Greek CO_2_ seep sites) for *C. nodosa* coincides with a range of seagrass species, for instance density and biomass increased with increase in CO_2_ levels, for *P. oceanica* (Hall-Spencer et al., 2008), *C. rotundata* and *C. serrulata* (Fabricius et al., 2011b; Russell et al., 2013) and *Zostera marina* (Palacios and Zimmerman, 2007), whereas decrease in biomass for *C. nodosa* in Vulcano CO_2_ seeps coincides with the results of Russell et al., (2013) for *Halophila ovalis* implying species-specific response to increased CO_2_ levels in future oceans. Higher density at all three CO_2_ seeps and higher horizontal rhizome biomass of *C. nodosa* at Vulcano among three CO_2_ seeps, suggest, *C. nodosa* at these seeps are adjusted to their carbon utilization capacity to the dissolved inorganic carbon (both CO_2 (aq)_ and HCO_3_^−^) concentrations of the surrounding water, similar results were obtained for *T. testudinum* in field and laboratory conditions (Durako., 1993). The low pH conditions were followed by increase in density (shoot and apex) of *C. nodosa* at all three CO_2_ seeps, compared to the reference sites, coinciding with results observed for *C. rotundata* and *C. serrulata* at Papua New Guinea CO_2_ seeps (Takahashi et al., 2016) and differing from previous results of *C. nodosa* at CO_2_ seeps of Vulcano (Apostolaki et al., 2014), whereas decrease of total biomass in Vulcano agrees with previous from the same area (Apostolaki et al., 2014). The above ground biomass was above 20% of total biomass at Greek and Italy CO_2_ seep sites, which agrees with findings for *C. nodosa* at Ischia seeps (Gianluigi et al., 2002), resulting in lower below ground biomass (lower than 80%) of seagrass meadows in elevated CO_2_ levels. Our results of lower below ground biomass did not match with results from another seagrass species (*Zostera marina*) in CO_2_ enriched conditions (Welsh et al. 1997; Palacios and Zimmerman, 2007).

Carbon content in *C.nodosa* leaves increased at the CO_2_ seep stations at Paleochori and Adamas sites, and remained invaribale at Vulcano sites, which agrees with experimental results from *T*. t*estudinum* and *T. hemprichii* which showed invaribale carbon content in CO_2_ enriched and reference conditions (Campbell and Fourqurean et al., 2013: Jiang et al., 2010). The N content of the *C. nodosa* leaves (1.67 % DW ± 0. 36) were low at Paleochori CO_2_ seep and were below the threshold (1.8% DW) for N limitation in seagrass (Duarte. 1990) suggesting that nitrogen deficiency may have limited overall plant growth, whereas N content of *C.nodosa* leaves at Adamas (1.85% DW ± 0.05) were not along the thresehold level and may have added to higher growth and density of plants at the Adamas CO_2_ seeps. In case of *C. nodosa* leaves at Vulcano even with higher N content, growth and density remained lower, similar results were obtained for *C. nodosa* in experimental CO_2_ and nutrient enrichment conditions (Khan et al., 2016). Simultaneously lower growth can be attributed to the epiphytic microalgae on seagrass leaves, which gets dominant during high N content, compete with the seagrass in production and eventually shade the seagrass leaves (Valiela et al., 1997). Lower C: N ratios in leaves of the plants at all sites (Table 3) suggests that plants near the seeps utilize the available nitrogen rapidly relative to their carbon content (Duarte. 1990). As nutrient availability increases seagrass tissues, must have been become progressively enriched in nitrogen relative to the carbon content implying decreasing C: N ratios, as found in rhizomes of all three CO_2_ seeps and roots of Vulcano seeps, whereas the opposite trends were observed in the roots of *C. nodosa* at Greek CO_2_ seeps (Table 3).

Significant interactions among stations and sites for *C. nodosa* variables (biomass, morphometry, carbon and nitrogen content) suggests that the effects of CO_2_ on these variables depends on the sites too, i.e., the levels of CO_2_ at these sites. This also indicates that positive response of biomass (total, horizontal and Ag: Bg), morphometry (no. of leaves) and carbon content (leaves and roots) to CO_2_ levels at the three sites are not entirely due to CO_2_ seeps but due to also the local environmental conditions of these seeps. Although there was an interaction among sites, ANOVA analysis (Tukey’s test) indicated which were the sites that have significant effect of CO_2_ on *C. nodosa* variables.

The low rhizome elongation (ca. 4.27 – 10.6 cm year^−1^) for *C. nodosa* at Adamas, Paleochori and Vulcano (ca.7.91 cm year^−1^) CO_2_ seeps suggested a reduced recruitment (Duarte and Sand-Jensen, 1990) compared to other populations elsewhere (40 cm year^−1^; Marbà and Duarte, 1998). Increased growth (vertical and horizontal elongation) and production (vertical rhizome, horizontal rhizome and leaf) with proximity to CO_2_ seeps at all three locations suggests plants were C limited in current conditions in the world oceans (Zimmerman et al., 1997; Invers et al., 2001). Additionally, the Adamas, Paleochori and Vulcano seeps, apart from CO_2_ supplies, may also be enriched volcanic inputs such as trace elements, which above certain levels can be toxic (Vizzini et al. 2013) and along with low pH and depleted oxygen levels can deteriorate the chemical properties of nearby water and sediment and in turn affect the seagrass productivity negatively.

Our results represent an early assessment of seagrass *C. nodosa* response under long term CO_2_ enrichment using population reconstruction techniques as a tool. Comparison to similar studies of *C. nodosa* at other seeps suggests that seagrass response to naturally acidified conditions is not so straight forward as the response is species-specific and depends on the biogeochemical characteristics of the site. Clearly, further research on seagrass growth and production using reconstruction techniques in CO_2_ seeps in world oceans is necessary, keeping in mind the ecosystem services seagrass provides, their capacity to act as global carbon sink and their major role in mitigation of the elevated levels of CO_2_ in future oceans.

## Acknowledgement

This work was part of MARES “Future Oceans” project (MARES _12_14) and was funded through a MARES Grant. MARES is a Joint Doctorate programme selected under Erasmus Mundus coordinated by Ghent University (FPA 2011-0016). Check <u>www.mares-eu.org</u> for extra information. We are grateful for the support of Thanos Dailianis, Julius Glampedakis and Veronica Santinelli in field sampling and laboratory works respectively at HCMR, Greece.

